# Senolytics and the compression of late-life mortality

**DOI:** 10.1101/2021.04.24.441236

**Authors:** Axel Kowald, Thomas B L Kirkwood

**Affiliations:** Ageing Research Laboratories, Campus for Ageing and Vitality, Newcastle University, Newcastle upon Tyne NE4 5PL, U.K.; Rostock University Medical Center, Institute for Biostatistics and Informatics in Medicine and Aging Research (IBIMA), Rostock, Germany; Center for Healthy Aging, Department of Cellular and Molecular Medicine, University of Copenhagen, Copenhagen 2200, Denmark

## Abstract

Senescent cells play an important role in mammalian ageing and in the etiology of age-related diseases. Treatment of mice with senolytics – drugs that selectively remove senescent cells – causes an extension of median lifespan but has little effect on maximum lifespan. Postponement of some mortality to later ages, without a corresponding increase in maximum mortality, can be termed ‘compression of mortality’. When we fit the standard Gompertz mortality model to the survival data following senolytic treatment, we find an increase in the slope parameter, commonly described as the ‘actuarial ageing rate’. These observations raise important questions about the actions of senolytic treatments and their effects on health and survival, which are not yet sufficiently understood. To explore how the survival data from senolytics experiments might be explained, we combine recent exploration of the evolutionary basis of cellular senescence with theoretical consideration of the molecular processes that might be involved. We perform numerical simulations of senescent cell accumulation and senolytic treatment in an ageing population. The simulations suggest that while senolytics diminish the burden of senescent cells, they may also impair the general repair capacity of the organism, leading to a faster accumulation post-treatment of new senescent cells. Our results suggest a framework to address the benefits and possible side effects of senolytic therapies, with the potential to aid the design of optimal treatment regimens.

## 1 Introduction

Ageing is thought to be driven by the gradual accumulation of molecular and cellular damage, causing progressive functional impairments (Kirkwood, 2005). A variety of processes including genomic instability, mitochondrial dysfunction, impaired proteostasis, and others have been categorized as ‘hallmarks of ageing’ (Kirkwood, 2005; Lopez-Otin *et al.*, 2013). Prominent among these is the phenomenon of cellular senescence, the defining feature of which is that cells enter a state of irreversible arrest. Many senescent cells also undergo alteration to produce the ‘senescence-associated secretory phenotype’ (SASP), involving the production of a complex array of chemokines, cytokines, growth factors and proteases, which cause significant effects on neighbouring cells, many of them apparently negative and including conversion into new senescent cells by way of the so-called ‘bystander’ effect (Nelson *et al.*, 2012; Xu *et al.*, 2017; da Silva *et al.*, 2018). In keeping with the idea that senescent cells are in some way harmful, recent experiments showed that in mice the targeted removal of senescent cells, termed ‘senolysis’, resulted in increased lifespan as well as beneficial effects on health (e.g. Baker *et al.*, 2011; Baker *et al.*, 2016; Baar *et al.*, 2017; de Keizer, 2017; Xu *et al.*, 2018; Yousefzadeh *et al.*, 2018).

Whilst work continues to explore the possible therapeutic benefits of senolysis, we recently suggested that it is important to ask what evolutionary forces might have been behind the emergence of cellular senescence (Kowald *et al.*, 2020). Entry into the senescent state appears to be regulated, presenting questions about why such a response should have evolved. It seems *a priori* unlikely that a purely negative action would be favoured by natural selection. In terms of potential benefits, cellular senescence is often regarded as an anti-cancer mechanism, since it limits the division potential of cells. However, many studies have shown that senescent cells often also have carcinogenic properties. Furthermore, other studies have shown that cellular senescence is beneficially involved in wound healing, development and tissue repair (Krizhanovsky *et al.*, 2008; Demaria *et al.*, 2014; Demaria *et al.*, 2015; Yun *et al.*, 2015; Ritschka *et al.*, 2017; Gibaja *et al.*, 2019). We recently brought these findings and ideas together and concluded that evolutionary logic strongly supports the idea that the latter positive contributions are the main reason for the evolution of cellular senescence. We further suggested that, since the immune system appears to play a role in clearing SASP-positive cells once they have performed their temporary functions, the observed age-related accumulations of senescent cells might arise simply because the immune system had to strike a balance between false negatives (overlooking some senescent cells) and false positives (destroying healthy body cells).

The importance of understanding the role of senescent cells is further indicated by recent senolysis studies in mice (Baker *et al.*, 2016; Xu *et al.*, 2018; Yousefzadeh *et al.*, 2018), where it was found that treatment with senolytics resulted in a substantial increase in mean and median survival times. However, in each of the studies there was much less increase in the maximum survival time. Such an outcome is only possible if, following senolytic treatments, the deaths that are postponed to produce the increased mean/median lifespans become concentrated in the interval prior to the relatively unaltered maximum lifespan. Such a phenomenon constitutes a ‘compression of mortality’, which needs to be explained. In this paper, we analyse in detail the changes in late-life mortality patterns and we then examine some ideas that might help to throw light on the mechanisms that are involved.

## 2 Analysis of mortality rates following treatment with senolytics

A statistical device commonly used to examine age-related mortality data is the Gompertz model (Gompertz, 1825) given by the equation

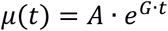

where *μ*(*t*) is the mortality rate at age *t*, *G* is a slope parameter (often termed the ‘intrinsic rate of ageing’), and *A* is the intercept in a logarithmic plot (often termed the ‘basal vulnerability’, i.e. mortality for t=0). The Gompertz model has found wide application in studies on ageing (Kirkwood, 2015) and has sometimes been described as a ‘law’ of mortality. In the present context, we avoid ascribing any deep biological meaning to the Gompertz model. After showing how the model fits the patterns of survival following treatment with senolytics, we simply use it as a means to describe and summarize these data.

We consider four studies that applied different types of senolytics to four different mouse populations and which published the relevant survival data (Baker *et al.*, 2016; Xu *et al.*, 2018; Yousefzadeh *et al.*, 2018). One way to analyse such data is to fit the Gompertz model to the logarithm of mortality, but unless this is based on a very large population size the calculation of mortality from survival data is quite error prone. To avoid these problems we fitted the Gompertz model directly to the survival curves, which are given by

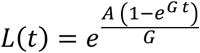

We pre-processed the raw data so that survivorship starts at the first day of treatment, normalised to 1. We used the function NonlinearModelFit of the software package Mathematica^®^ to estimate values for the parameters *A* and *G*. The function also provides confidence intervals and an overall measure for the goodness of fit (r^2^). In Fig. 1 the survival data are shown together with the survivorship curves. All fits result in r^2^ values > 0.99, indicating that the Gompertz model fits the post-treatment mortality data very well. In the following, we will now describe in more detail the four different experiments.

**Fig. 1:**
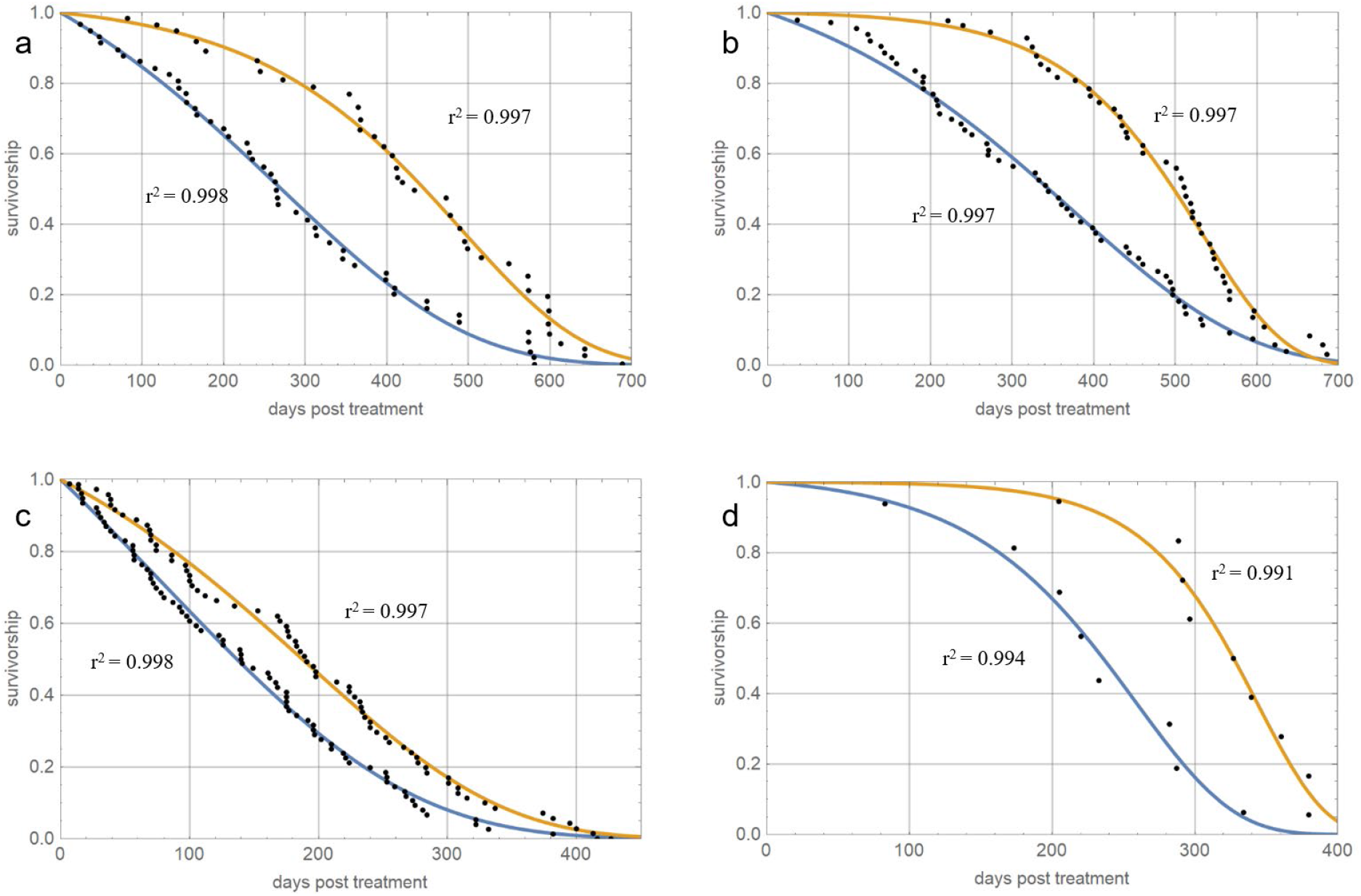
**(a)** Survival data for mice with mixed genetic background starting at the beginning of senolytic treatment (digitized from Baker et al. (2016)) overlaid with survival curves based on fitted Gompertz parameters (r^2^=0.997). **(b)** Survival data for mice with a C57BL/6 background starting at the beginning of senolytic treatment (digitized from Baker et al. (2016)) overlaid with survival curves based on fitted Gompertz parameters (r2=0.997). **(c)** Post-treatment survival data for mice treated with a senolytic cocktail of D&Q (data from Xu et al. (2018)) overlaid with survival curves based on fitted Gompertz parameters (r^2^=0.998). **(d)** Post-treatment survival data for mice treated with the senolytic compound fisetin (data from Yousefzadeh et al. (2018)) overlaid with survival curves based on fitted Gompertz parameters (r^2^=0.993).

Baker *et al.* (2016) investigated the effects on the lifespan of wild-type mice of removing cells identified as senescent by being positive for p16, a recognized marker of senescence. They treated two populations of genetically modified mice with different genetic backgrounds (mixed and C57BL/6) with the drug AP20187, which induces apoptosis in p16 expressing cells. The treatment was started at one year of age and injections were given twice weekly until the end of life. In the mice with mixed genetic background (59 treated animals, 57 controls), median lifespan was increased by 27% (from 624d to 793d). By contrast, maximum lifespan (using the 90% percentile as a proxy) was increased only by ca. 2.8%. As mentioned above, the fit of the Gompertz model resulted in an excellent agreement with the data (r^2^=0.998 for control and 0.997 for treatment group) as can be seen in Fig. 1a. The estimates for the Gompertz parameters *G* and *A* (with 95% confidence intervals) are shown in Fig. 2a. As can be seen from the non-overlapping confidence intervals, the rate parameter *G* is significantly higher in the treatment group than in the controls, while the intercept parameter *A* is lower.

**Fig. 2:**
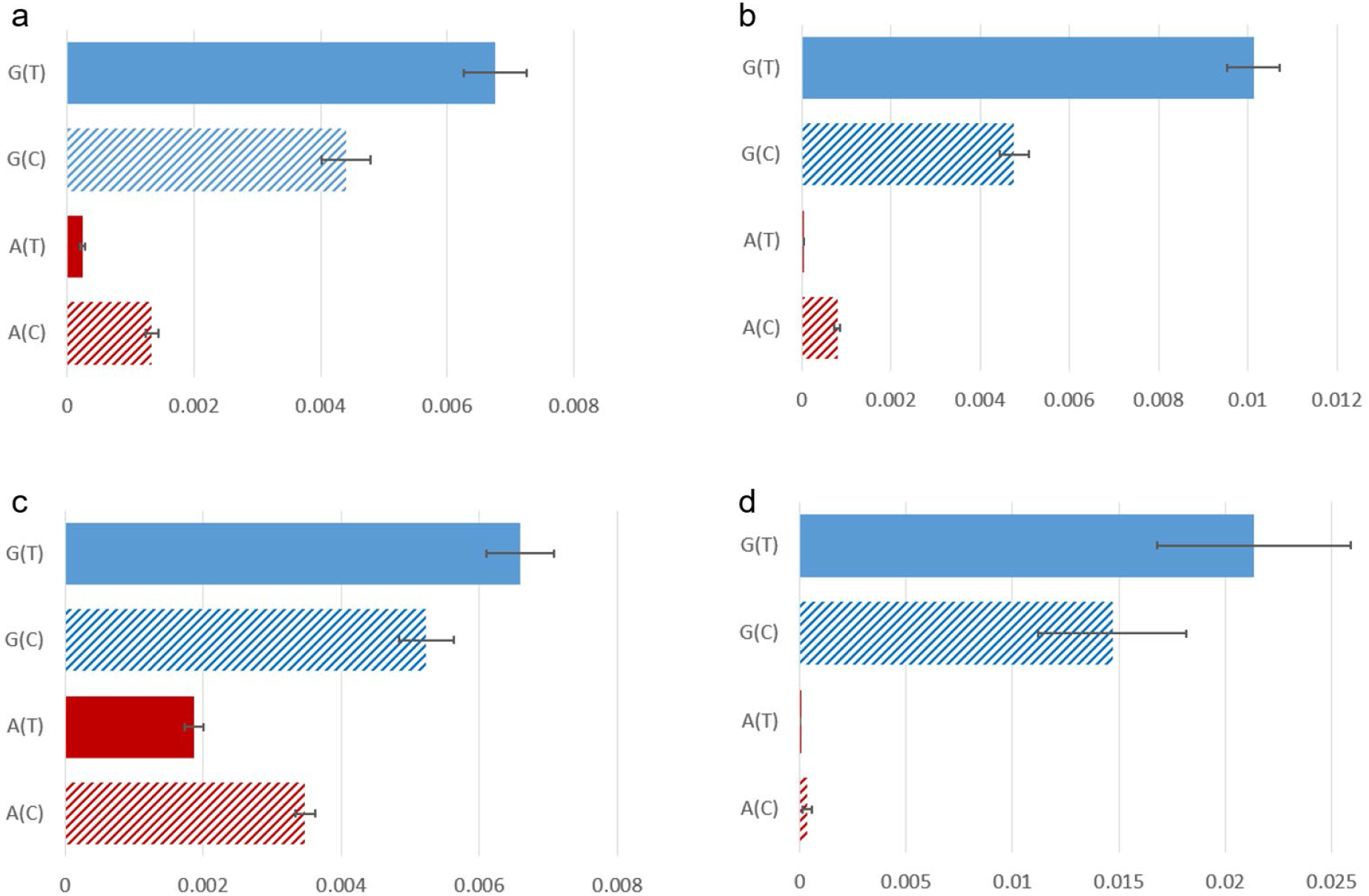
Estimates (with 95% CI) of Gompertz parameters G (blue) and A (red) for the treatment (T) and control (C) group of (a) Baker et al. (2016) (mixed genetic background), (b) Baker et al. (2016) (C57BL/6 genetic background), (c) Xu et al. (2018), and (d) Yousefzadeh et al. (2018).

In their mice with a C57BL/6 background (51 treated animals, 58 controls), Baker *et al.* (2016) found that median lifespan was increased by 24% (from 699d to 866d) while maximum lifespan (using the 90% percentile as a proxy) was only increased by ca. 10.5%. Applying the Gompertz model again resulted in an excellent fit (r^2^=0.997), as can be seen in Fig. 1b. The estimates for the relevant Gompertz parameters are shown in Fig. 2b. Also in this case, the rate parameter *G* is significantly higher in the treatment group than in the controls, while the parameter *A* is lower.

A further study was reported by Xu *et al.* (2018), who investigated the effects of a senolytic cocktail of dasatinib plus quercetin to eliminate senescent cells from a population of C57BL/6 mice. Treatment began at 24-27 months of age (71 treated animals, 76 controls) and was repeated biweekly until the end of life. Median lifespan post-treatment was significantly increased by 36% (from 140d to 191d), which corresponds to an absolute increase in median lifespan of 6.3% (from 937 to 996d). Maximum lifespan (using the 90% percentile as proxy) was increased by 19.6% post-treatment and by 5.3% over the whole life. Applying the Gompertz model once more resulted in an excellent fit (r^2^=0.998 for controls and 0.997 for the treatment group) as can be seen in Fig. 1c. The estimates for the relevant Gompertz parameters are shown in Fig. 2c. Again, the rate parameter *G* is significantly higher in the treatment group than in the controls, while the parameter *A* is lower.

Lastly, Yousefzadeh *et al.* (2018) investigated the senolytic compound fisetin using a smaller population of WT f1 C57BL/6:FVB mice starting at 85 weeks of age (9 treated animals, 8 controls). Fisetin was administered in the diet from the beginning of treatment until the end of life. Since the population size is in this case small, we calculated lifespan statistics from the fitted Gompertz curves instead of the raw data. Median lifespan post-treatment was increased by 38.7% (from 236d to 327d), which corresponds to an absolute increase in median lifespan of 11.0% (from 831d to 922d).

Maximum lifespan (using the 90% percentile as proxy) was increased by 21.3% post-treatment and by 7.4% over the whole life. Quality of fit was good (r^2^=0.994 for controls and 0.991 for the treatment group), but less than in the other studies (Fig. 1d). The estimates for the Gompertz parameters are shown in Fig. 2d. The estimates for *G* and *A* show the same tendency as in the other studies, but because of the very small sample size, the differences between treatment and control group do not reach statistical significance (i.e. the confidence intervals overlap).

To our knowledge, the above are all of the currently published studies that investigated lifespan effects of senolytic treatments. A recent online conference presentation (www.longevity2020.com) by OISIN Biotechnologies described use of their proprietary lipid nanoparticle (LNP) technique to deliver plasmids into cells that have a caspase behind a p16 or a p53 promoter. Thus, cells that express either of these genes are driven into apoptosis. The lipid nanoparticles were injected monthly into mice, starting at 106 weeks of age (9 treated animals, 12 controls). Once more, median lifespan was extended while maximum lifespan remained almost unchanged. A Gompertz fit indicated that in this case too, the treatment increased G while reducing parameter A (unpublished data kindly provided by OISIN). As in case of Yousefzadeh *et al.* (2018), the smallness of sample size did not admit statistical significance.

It might be noted that the estimates of the *A* parameter for the study of Xu *et al.* (2018) (Fig. 2c) are higher than for the other studies. The reason is most likely that in this investigation the mice had the highest age when treatment started (24-27 months). Since the parameter estimation was performed for ‘days post treatment’, parameter *A* represents mortality at the beginning of treatment and not at birth, so the estimates for *A* should indeed be highest for this study. The estimation of parameter *G*, however, should not be affected by the timing of the treatment since the slope of the mortality increase is more or less independent of age. Accordingly estimates for *G* are comparable for all studies except for Yousefzadeh *et al.* (2018), where they are higher for controls as well as the treatment group. The explanation here might be that the mouse strain used by the authors has a shorter lifespan (tMedian = 830 days) than for instance the mice used by Xu *et al.* (2018), (tMedian = 937 days). Consequently, that strain could exhibit a higher value for *G*. However, independently of these considerations, the important observation is that in all studies a senolytic treatment always led to a drop of parameter *A* but an increase of parameter *G*.

## 3 How might senolysis result in the observed lifespan effects?

We will now use simple mathematical models to explore how senolysis might act upon age-related molecular and cellular damage in such a way that we can reproduce the effects observed in the lifespan studies. Since experimental data to explain these effects are not yet available, we will use the models for exploratory and illustrative purposes. To formulate the models, we consider a number of hypothetical scenarios. Each scenario examines possible ways in which damage might accumulate and the consequences of applying senolytics. With ageing, accumulation of damage occurs that results in an increasing mortality rate. The damage can be of several kinds, reflecting the diversity of mechanisms that result in the various hallmarks.

**Scenario A** supposes simply that all damage is associated with cellular senescence and its adverse effects on health. From some initial level, *D*(0), damage accumulates proportional to the product of the existing amount of damage and a rate parameter *c*. The equation relating damage level at time *t*+1 to the level at time *t* is written

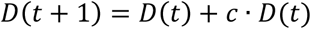

When a senolytic treatment is applied, the existing damage level is immediately reduced by some fraction *hD*, thereafter damage continues to accumulate again, starting from the reduced level. For simplicity we assume that damage is equivalent to mortality risk. Although it seems unlikely that all varieties of damage will be reduced by senolytics, this first scenario establishes a baseline.

**Scenario B** supposes a similar mechanism for damage accumulation but now includes two classes of damage. Class 1 represents damage in which cellular senescence plays the central role. In this class, senolytics will have a direct effect of reducing the level of accumulated damage, as in scenario A. Class 2 represents damage in which cellular senescence *per se* plays no role, but this scenario will explore the possibility of cross-talk between senescence-related and senescence-unrelated damage. Equations relating damage level in each class at time *t*+1 to the level at time *t* are written

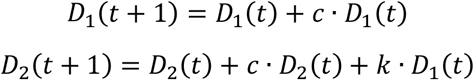

If parameter *k* is zero, then damage in class 2 is completely independent of senescent cells and senolytics. However, if *k* is positive it represents indirect actions of senescent cells that might affect class 2. Effects of senescent cells on Alzheimer (Zhang *et al.*, 2019), atherosclerosis, fibrosis or cancerogenesis (Yanai & Fraifeld, 2018) would be such examples. In such circumstances extra survival benefit can be expected from senolysis, since it would indirectly also affect class 2 damage.

Finally, in **Scenario C**, we consider the possibility that senolytic treatments might interfere with the machinery that normally acts to remove senescent cells. It is known that some cells express p16 without being senescent, an example being macrophages (Hall *et al.*, 2017). Since macrophages and other components of the immune system are involved in the removal of senescent cells (Yun *et al.*, 2015; Burton & Stolzing, 2018; Ovadya *et al.*, 2018), senolytic drugs might temporarily interfere with this process until they are cleared from the body. We model this idea by adding a further variable, *RC*, to denote the ‘repair capacity’, which might be regarded as a high-level representation of the immune system. With this the equations become

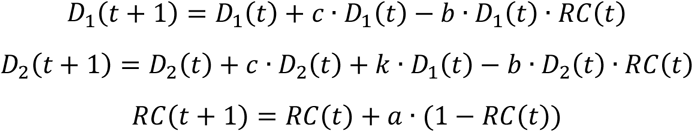

The damage of class 1 (representing senescent cells) is now diminished by a further term that is proportional to the existing damage and the current repair capacity. The new equation for *RC* is constructed in such a way that it approaches a steady state of one caused by a combination of constant production rate and degradation at a rate proportional to the existing amount. The parameter *a* controls how fast the equilibrium is regained after a disturbance. In this scenario, a senolytic treatment would instantaneously decrease *D*_1_ as well as *RC* (although by different fractions *hD* and *hRC*).

## 4 Model structure and simulation methods

To investigate the behaviour and properties of the different model scenarios we developed software using Python that calculates survival curves of a population of 100,000 individuals based on specific parameter values. The software creates new-born individuals with an initial amount of damage, *D*(0), assigned to damage class 1 (Scenario A) and class 2 (Scenarios B and C). Although the scenarios only differentiate between two major classes of damage, it is clear that in reality *D*_1_ and *D*_2_ consist of many subclasses that represent different types of age-related damage (defective mitochondria, protein cross-links, etc.). We deal with this situation by creating a total of 100 subclasses of damage that can either be assigned to damage class 1 or class 2. This provides a further model parameter, *f*, that controls which fraction of the overall age-related damage responds to senolytic treatment (*D*_1_) or not (*D*_2_). The program then loops through the life course in time intervals of one week and performs the following computational steps:

- For each individual the amounts of damage class 1 and class 2 (represented in terms of each one of their respective subclasses), as well as the repair capacity (*RC*), are updated according to the equations and parameters of the corresponding scenario.
- For each individual a random number is generated and based on the current level of damage (i.e. mortality risk) of all different subclasses it is decided if the individual died or not. If yes, the individual is removed from the population.
- The program checks if at this time point a senolytic treatment should be applied. If yes, for each individual the amount of damage in all subclasses belonging to *D*_1_ is reduced by *hD*. If we are in Scenario C, the current level of the repair capacity is also reduced by *hRC*.
- If there are individuals still alive, the loop is repeated; otherwise, the simulation is terminated and the results are recorded.

## 5 Results of simulations

### 5.1 Scenario A

Fig. 3 shows the simulation results for a cohort without any interventions. Median lifespan is reached after 125 weeks (Fig. 3a) and mortality increases linearly on a logarithmic scale (Fig. 3b), starting from damage (i.e. mortality) *D*(0) for *t*=0. For very young and very old ages, the mortality curve fluctuates more strongly because of either very few deaths occurring or very few individuals being alive. This simulation serves as reference for the following simulations with senolytic intervention.

**Fig. 3:**
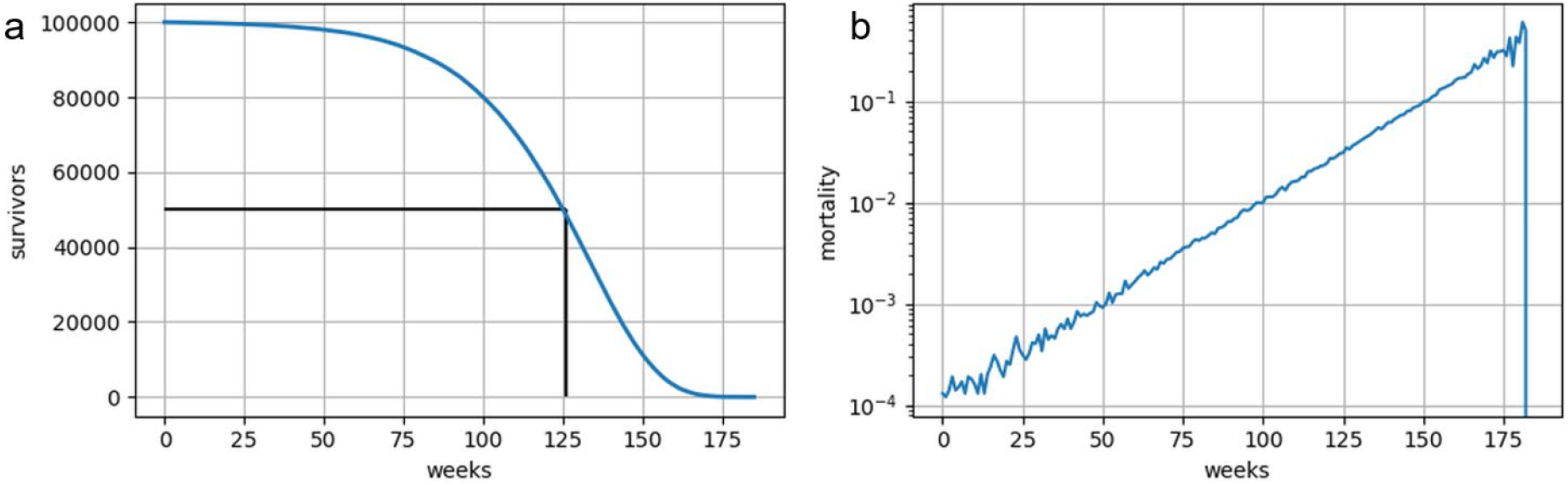
Life course simulation without senolytics intervention. Parameters *D*(0) and *c* are chosen to reproduce the survival curve of a typical wt mouse with a median lifespan of 125 weeks (indicated by black lines in (a)) and a max*imum* lifespan of 166 weeks (*D*(0) = 0.0001141 week^−1^, *c* = 0.04658 week^−1^). The figure shows survivorship (a) as well as mortality (b) per week over time.

To understand how a single senolytic treatment might influence the life course of a cohort we performed a simulation with a senolytic treatment at week 80 that reduces damage by 80% at that time point (*hD*=0.8). We chose this limited efficiency since it is known that senolytic compounds do not remove all senescent cells. As can be seen from Fig. 4a, this treatment has a strong effect on the median as well as maximum lifespan. The former is increased by 27% (from 125 to 159 weeks) and the latter by 20% (from 166 to 202 weeks). At the time point of treatment, mortality drops abruptly (since the damage causing mortality is reduced) but then rises again, as new damage accumulates.

**Fig. 4:**
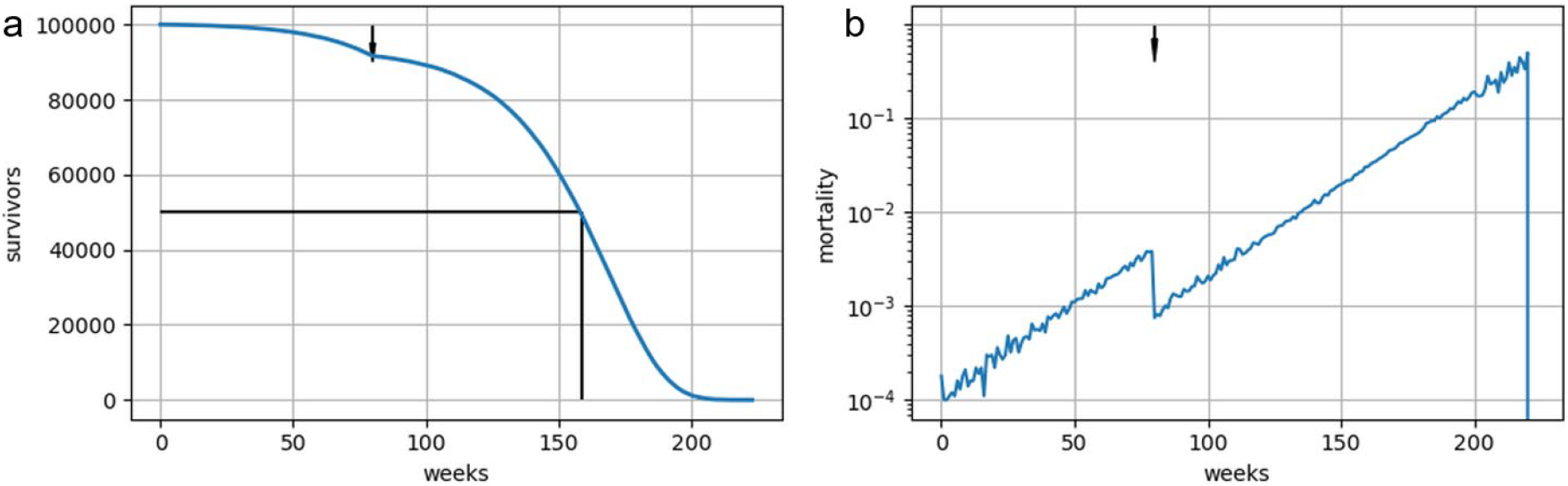
Life course simulation with a single senolytics intervention at 80 weeks of age (indicated by arrow). The reduction of damage, *hD*, is set to 80%, which results in a steep drop of mortality at the time of treatment (b). As a consequence, median lifespan is increased by 27% and maximum lifespan is increased by 20% compared to the situation without treatment. The other parameters are as in Fig. 3.

However, in each of the experimental studies we have described, the senolytic treatment was given not once but repeatedly after the start of the therapy. Simulating such a treatment regimen in Scenario A is possible, but also shows its limitations. Repeated reduction of all age-related damage effectively eliminates damage and thus mortality, which in turn leads to a non-ageing population with practically unlimited lifespan (simulation not shown). This is at odds with the real experimental observations and thus it is necessary to move on to less simplistic scenarios.

### 5.2 Scenario B

This scenario introduces a second damage class (*D*_2_), unaffected by senolytic treatment, as well as a cross-talk parameter, *k*, by which damage of class 1 can cause additional damage of class 2.

We investigate Scenario B by simulating a single senolytic treatment at week 80 with a small value for *k*. As can be seen in Fig. 5c the two different classes of damage behave quite differently. *D*_1_, which is susceptible to senolytics drops abruptly upon treatment (red), while *D*_2_ is not affected at all (blue). In this simulation 80% of all age-related damage was assigned to *D*_1_, which is reflected by the fact that at birth most of the damage belongs to this class. The cross-talk parameter, however, causes an accelerated accumulation of damage *D*_2_ so that around the time of treatment damage in both classes has accumulated to comparable quantities. This explains why the drop in mortality is so much smaller than in case of Scenario A (Fig. 4). Consequently, median lifespan did not change and maximum lifespan was even diminished by 1% through the senolytic treatment.

**Fig. 5.**
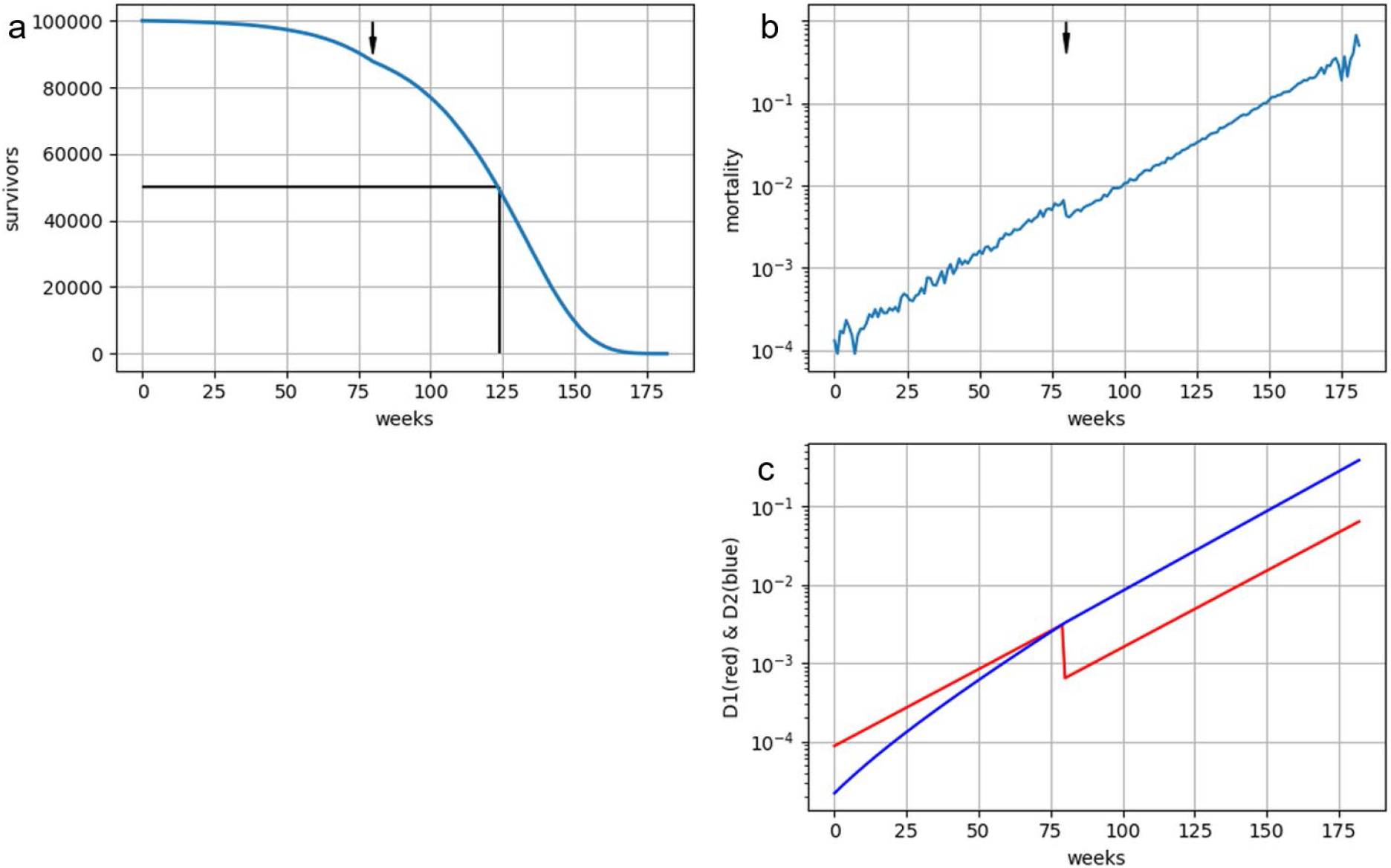
(a & b) Life course simulation with a single senolytic treatment at week 80 (indicated by arrow) according to Scenario B. (c) 80% of damage classes belong to class 1 (red), which responds to senolytic treatment by a reduction of damage (*hD*=0.8). The cross-talk parameter, *k*=0.01, strongly accelerates the accumulation of damage of class 2 (blue), so that the senolytic effects on lifespan are basically abolished (median +0%, max - 1%). The other parameters are as in Fig. 4.

To study the effect of the cross-talk parameter in greater depth we performed simulations with different values of *k* and plotted the resulting changes of median and maximum lifespan in Fig. 6. As can be seen, the results of Scenario B are very sensitive to *k*. Even small values of *k* lead to such an aggressive accumulation of senolytics resistant damage (*D*_2_) that a senolytic treatment is practically non-effective.

**Fig. 6.**
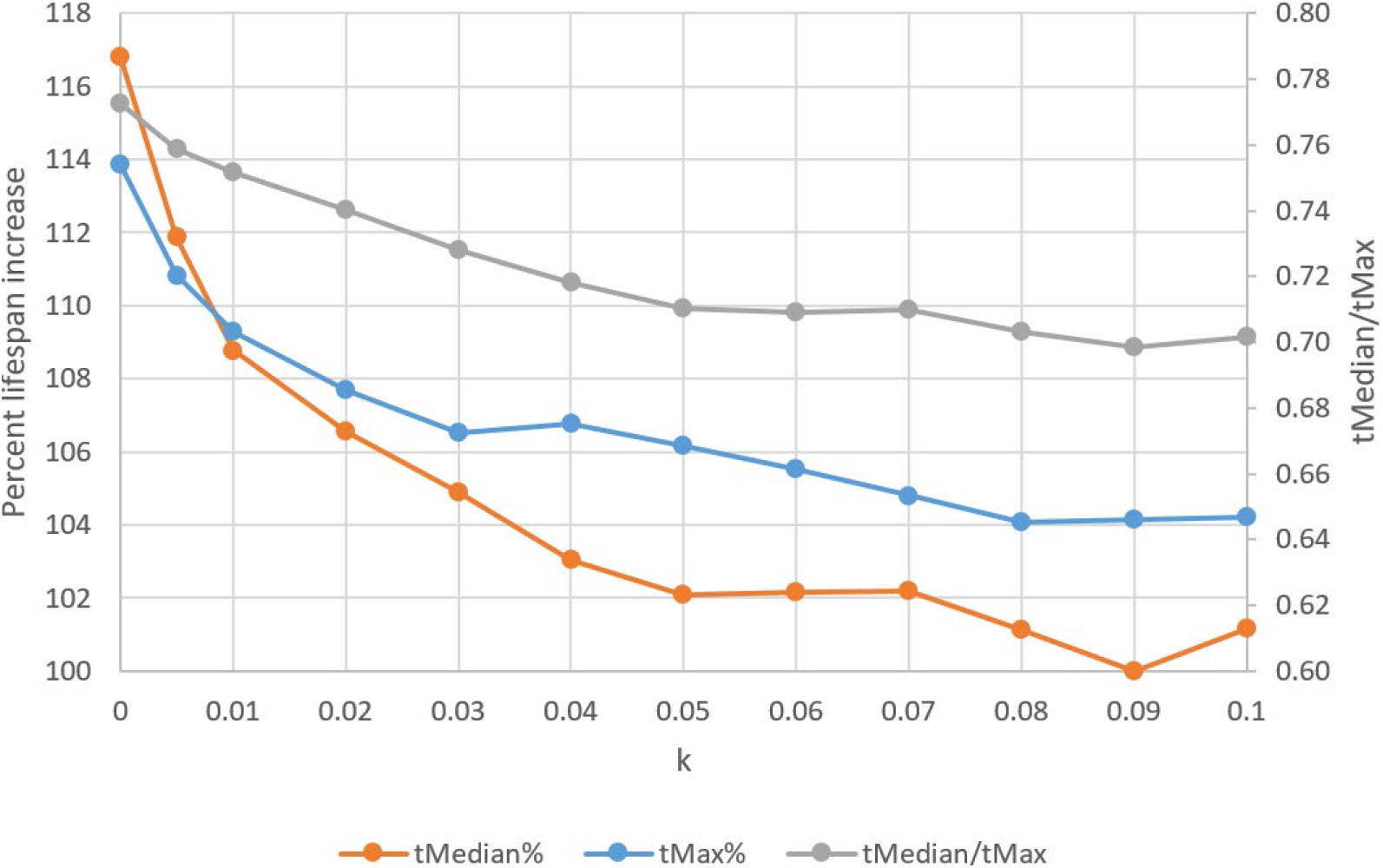
Variation of the cross-talk parameter, *k*, with a single dose of senolytics at 80 weeks of age. The resulting lifespan extension relative to the case without senolytics is shown on the left axis, while the ratio of tMedian to tMax, a measure of compression of mortality, is shown on the right axis. Even small values of *k* accelerate the accumulation of class *2* damage so strongly that the senolytic treatment is rendered ineffective and essentially abolishes lifespan extension. Other parameters are as used for Fig. 5

### 5.3 Scenario C

Senolytic substances do not have perfect specificity. Since senescent cells are quite heterogeneous and lack a universal marker, which would allow their identification, all the senolytic drugs currently available display not only a false negative but also a false positive rate. As a consequence of this, as mentioned earlier, senolytic treatments might also impair components of the damage repair system. Scenario C takes this into account, and Fig. 7c shows how a single senolytic treatment at 80 weeks of age temporarily reduces the capacity, *RC*, of the repair system. For the simulation it was assumed that *RC* is reduced by 10% through the treatment and recovers with a rate controlled by parameter *a*. While *RC* is impaired by the senolytic treatment, the removal rate of senescent cells is reduced, resulting in a faster net accumulation rate of all age-related damage. This effect, however, can hardly be seen in Fig. 7d since recovery of *RC* is fast and *hRC* is small. Consequently, median lifespan is increased by 14% and maximum lifespan by 11% compared to no treatment.

**Fig. 7.**
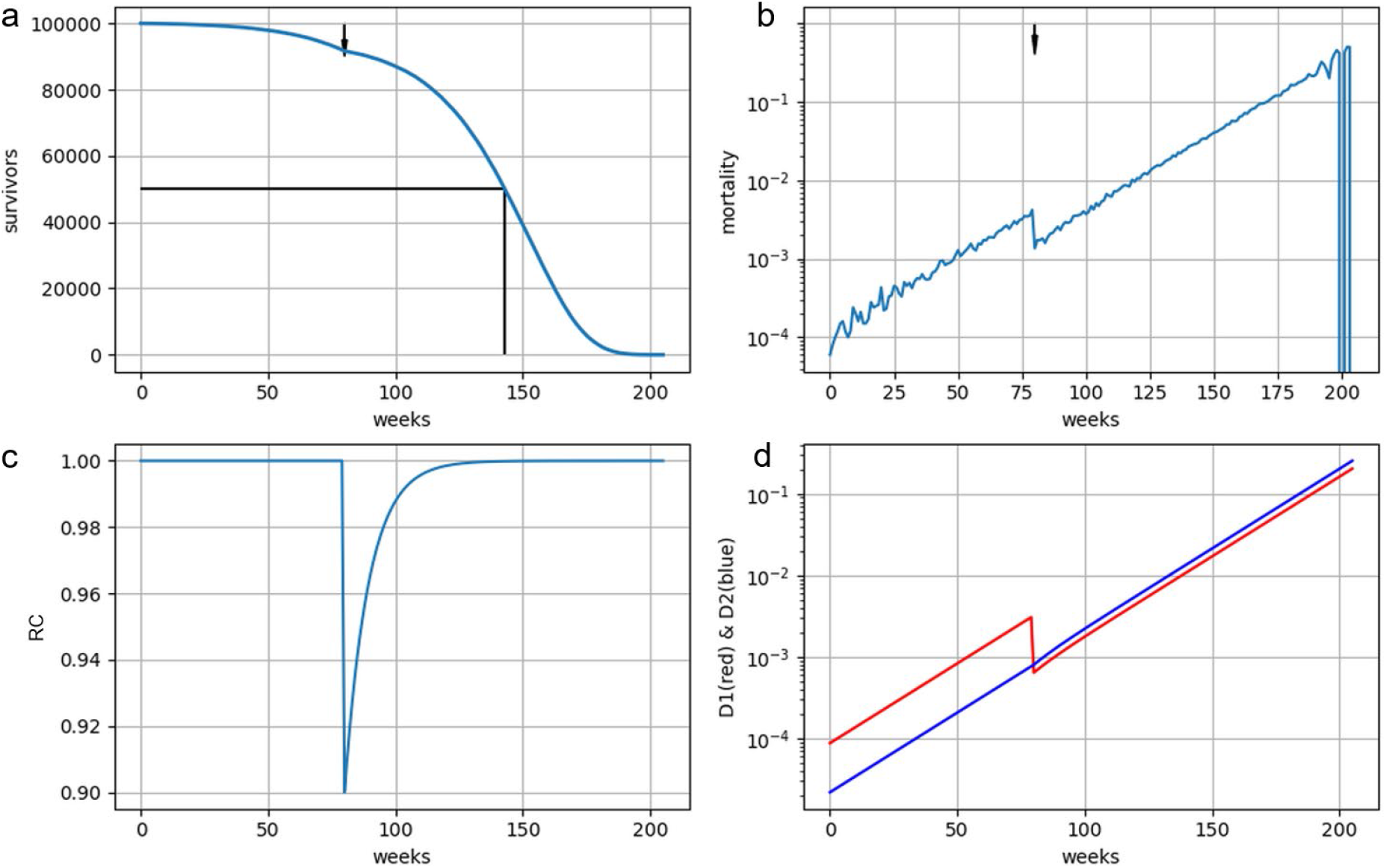
(a & b) Life course simulation with a single senolytic treatment at week 80 (indicated by arrow) according to Scenario C. 80% of damage subclasses belong to class 1 (red), which responds to senolytic treatment by a reduction of damage of 80% (*hD*=0.8) (d). Damage of class 1 is reduced by the repair system (*b*=0.154), but senolytic treatment also reduces repair *capacity* by 10% (*hRC*=0.1), shown in panel (c). RC recovers to its steady state value, with a rate given by parameter *a* (here 0.1). The rate of damage creation was adjusted to *c*=0.2, so that without senolytic treatment it results, in combination with the damage clearing effect of the repair system, in the same survival curve as for scenarios A and B.

The situation changes significantly, however, if we include repeated treatments. Baker *et al.* (2016) and Xu *et al.* (2018) administered their drug bi-weekly, while Yousefzadeh *et al.* (2018) added it to the diet until death, so that the animals received it continuously.

Fig. 8 shows the simulation results when treatment was started at 80 weeks and then repeated every 4 weeks. Although we assumed only a modest reduction of the repair system (*hRC*=0.1) the repeated treatment led to a steady state reduction of ca. 22% (from 1 down to 0.78) (Fig. 8c). Damage classes *D*_1_ and *D*_2_ react quite differently to the repeated schedule. While *D*_1_ damage is permanently repressed to a negligible level, *D*_2_ damage actually accumulates faster after start of the treatment (Fig. 8d). This is caused by the fact that clearance of age-related damage via the repair system *RC* is now reduced by the above-mentioned 22%. These processes, going on at the level of molecular and cellular damage, then manifest themselves phenotypically through a higher rate of mortality increase (Fig. 8b), which appears as an increase of Gompertz parameter *G* in the analysis of the experimental studies (Fig. 2). Consequently, in this simulation repeated senolytic treatment increased median lifespan more strongly (+9%) than maximum lifespan (−1%).

**Fig. 8.**
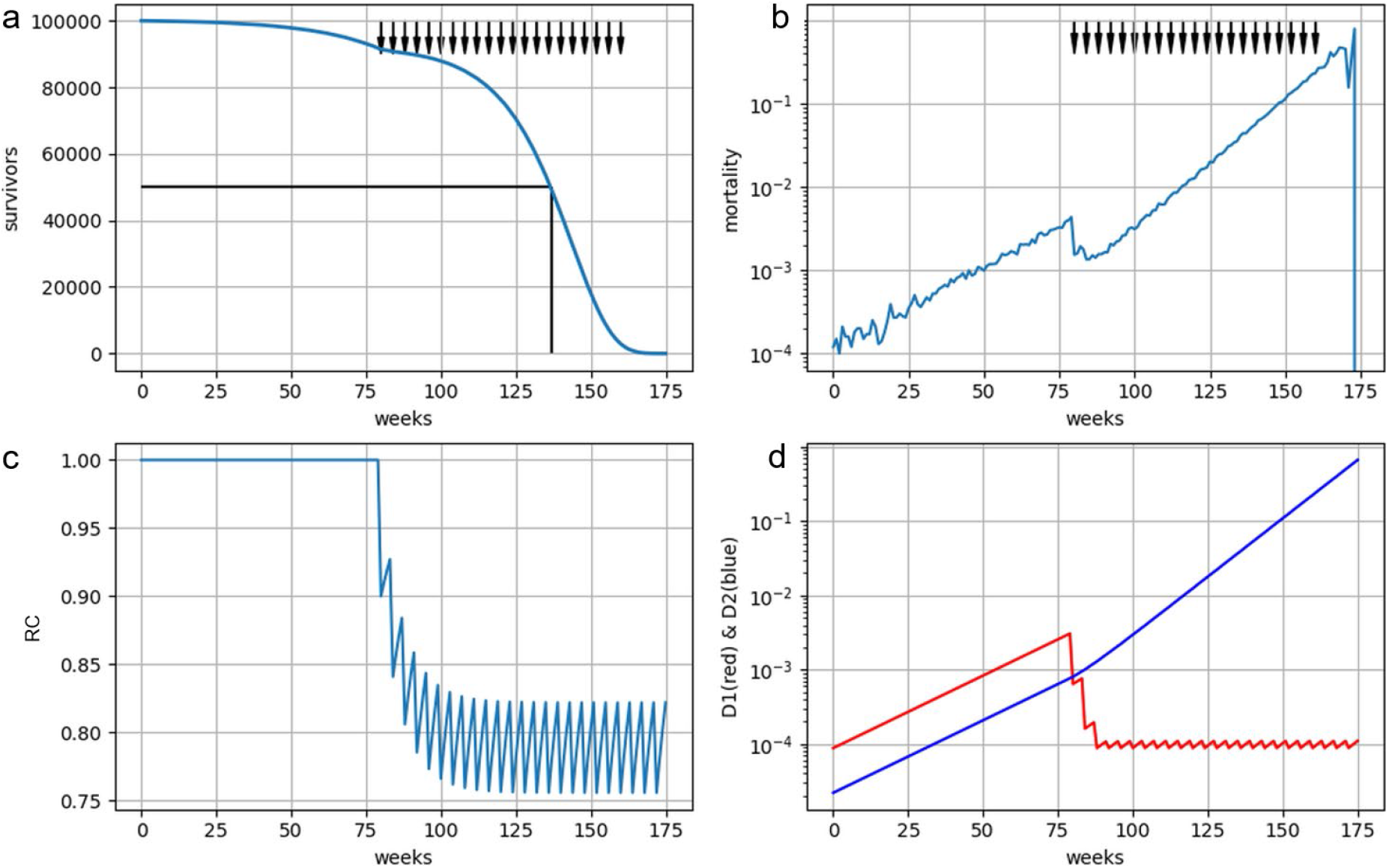
(a & b) Life course simulation of Scenario C with senolytic treatment starting at week 80 and repeated every 4 weeks (indicated by arrows), which each time reduces the amount of damage in 80% of damage classes by 80% but also reduces *RC* (*hRC*=0.1). As a consequence class 1 damage is permanently reduced to an extremely low level (we assume that it cannot be reduced below its *D*(0) value) while class 2 damage now accumulates faster than before, caused by the permanent impairment of the repair system.

### 5.4 Optimized treatment schedule

An important insight from Scenario C is that repeated treatments with senolytic compounds can have both desired effects (removal of susceptible damage *D*_1_) and undesired effects (accelerated accumulation of non-susceptible damage *D*_2_) (Fig. 8). This could provide a possible explanation for the experimental observation that senolytic treatments extend maximum lifespan much less than median lifespan. Under this assumption, an optimized treatment schedule might be to have only a single treatment since this only temporarily affects the repair capacity.

For the simulation shown in Fig. 7 the age of treatment, senoTime, was selected to be 80 weeks. This, however, is a parameter that can be chosen freely by the experimenter. Fig. 9 displays the consequences for median and maximum lifespan if senoTime is varied from 40 to 160 years. As can be seen, the most effective strategy is to perform the treatment as early as possible. For early treatment ages the relative gain in maximum lifespan is smaller than the gain of median lifespan, but this changes as senoTime is increased, since median lifespan extension must drop to 100% as senoTime approaches the median lifespan without treatment. The different influence of senoTime on median and maximum lifespan also means that the ratio of median to maximum lifespan, which we can interpret as a measure of compression of mortality, varies with the age of treatment. If a high value is desirable (see discussion) then an early treatment age should be chosen.

**Fig. 9.**
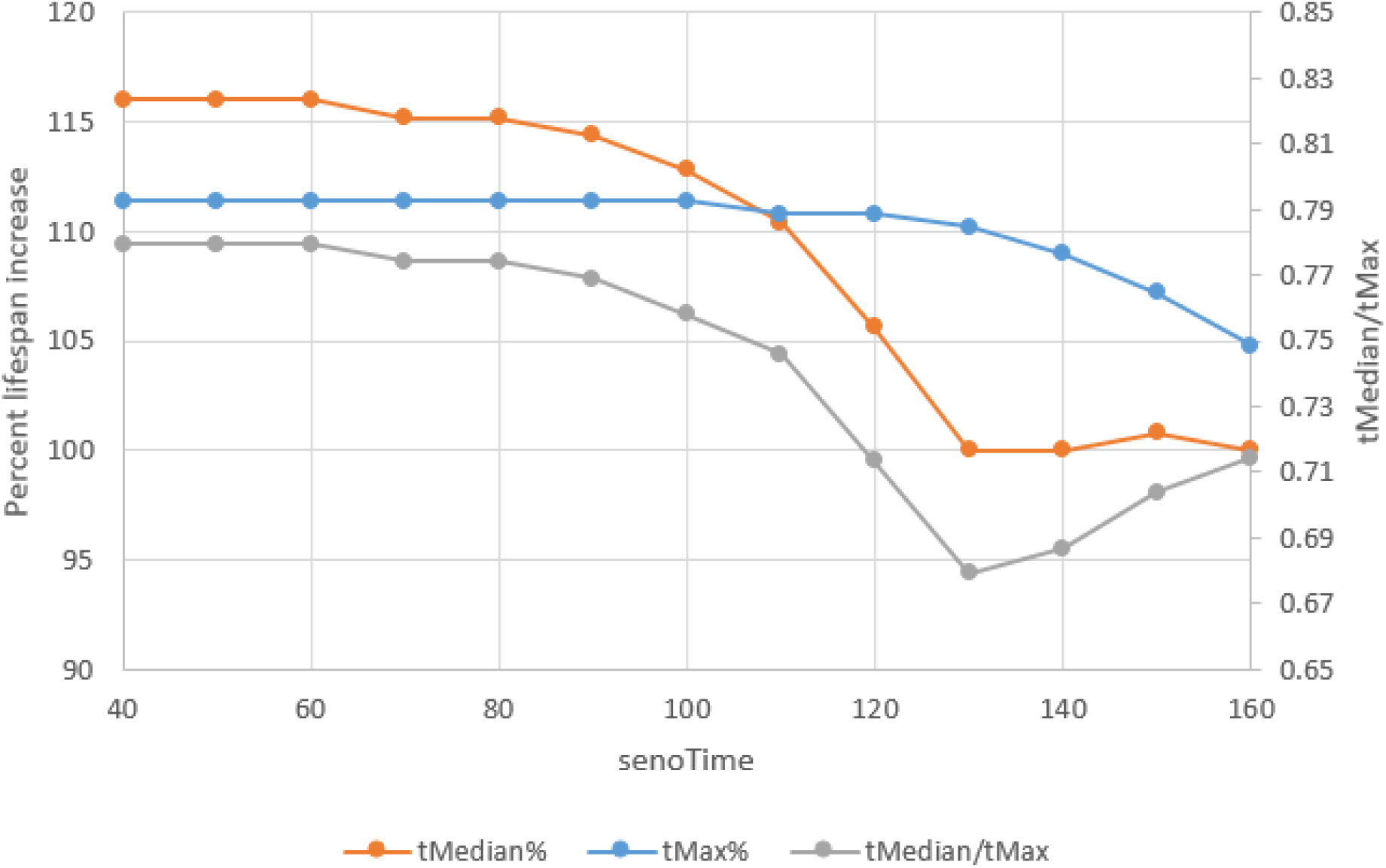
Variation of the time of treatment, senoTime, with a single dose of senolytics according to Scenario C. The resulting relative lifespan extension is shown on the left axis, while the ratio of tMedian to tMax, a measure of compression of mortality, is shown on the right axis. While repeated treatments only minimally exten*d* maximum lifespan (Fig. 8), a single treatment does not show this behaviour. If treatment time is beyond median lifespan, tMedian cannot be extended, but the treatment still increases maximum lifespan. Other parameters are as used for Fig. 8

## 6 Discussion

In the first part of this paper, we performed a statistical analysis of post-treatment survival in the four experimental studies in mice. In each study, senolytic treatment led to a substantial increase in median lifespan, while the effect on maximum lifespan was much smaller. The data showed an excellent fit to the Gompertz logarithmic equation, thus allowing description in terms of two summary statistics: the intercept parameter *A* and the slope parameter *G*. Note that in performing this analysis, no assumptions are made; the findings are a statement of statistical fact.

Table 1 summarizes these findings. For each experiment, the table shows the age in weeks at which treatment was started. The next two columns show the values of *A*_TC_ and *G*_TC_, which are the values of *A* and *G* for the treated animals as a percentage of the *A* and *G* values of the control animals. The final two columns show median (tMedian_TC_) and maximum (tMax_TC_) lifespans, again in percentage terms for treatment versus control. There is broad consistency in the pattern of results across all experiments although, as already remarked in section 2, there are some differences in the *A*_TC_ and *G*_TC_ values for Xu *et al.* (2018), which most likely relate to the later start of treatment in this study.

**Table 1.** Summary of the life extending effects (given as ratio between the treatment and control group*s*) in the four senolytic studies including the effects seen on the Gompertz parameters *A* and *G* (see also Fig. 2). The time of starting *t*reatment is given in weeks.

| Study | Treatment start | $A_{TC}$ | $G_{TC}$ | tMedian<sub>TC</sub> | tMax<sub>TC</sub> |
| --- | --- | --- | --- | --- | --- |
| Baker et al. (2016)<br>(mixed genetic background) | 52 | 18% | 154% | 27% | 2.8% |
| Baker et al. (2016)<br>(C57BL/6 genetic background) | 52 | 6% | 213% | 24% | 10.5% |
| Xu et al. (2018)<br>(C57BL/6) | 104-116 | 54% | 126% | 6% | 5.3% |
| Yousefzadeh et al. (2018)<br>(WT f1 C57BL/6:FVB) | 91 | 4% | 145% | 11% | 7.4% |

The increase in the value of *G* following senolytic treatments is a particularly striking observation, which we are not aware has been previously remarked. It requires explanation.

In the second part of the paper, we developed computer simulations of three possible mechanistic scenarios in order to gain a better understanding of possible modes of action of senolytic treatments. Scenario A, which supposes simply that senescent cells are all-important in ageing, was shown to be incompatible with experimental findings. Scenario B, which allows for other forms of damage to be involved and which also allows for senescent cells to drive these other forms of damage to some degree, was also found not to explain the data, although it does generate some interesting behaviours. In contrast, Scenario C proved to have the potential to explain the experimental findings. Scenario C includes the idea that the immune system plays an important role in removing senescent cells and related damage, but that this ‘repair capacity’ of the immune system is also negatively affected by senolytic drugs. In the case of a single senolytic treatment the repair capacity can recover (Fig. 7), but if the treatment is given continuously (as in all the experimental studies), the repair capacity is chronically reduced. This leads to an accelerated accumulation of damage, causing a faster increase of mortality, i.e. higher *G* value (Fig. 8).

Can the mechanism represented in Scenario C explain the experimentally observed changes in *A* and *G*? Table 1 shows that *A*_TC_ can be as low as 4% and *G*_TC_ as high as 213%. From the equations for Scenario C it can be seen that the phenotypically observable parameter *G* is given by *c*-*b* (if *RC* is at its steady state value of 1). Repeated senolytic treatments reduce the level of *RC*, resulting in a value of *G*_TC_= (*c*-*b*\**RC*)/(*c*-*b*). Thus, if we assume that roughly 75% of damage is removed by *RC* (*c*=0.2 and *b*=0.154), already a modest reduction of *RC* can explain the measured values for *G*_TC_ (Fig. 10).

**Fig. 10.**
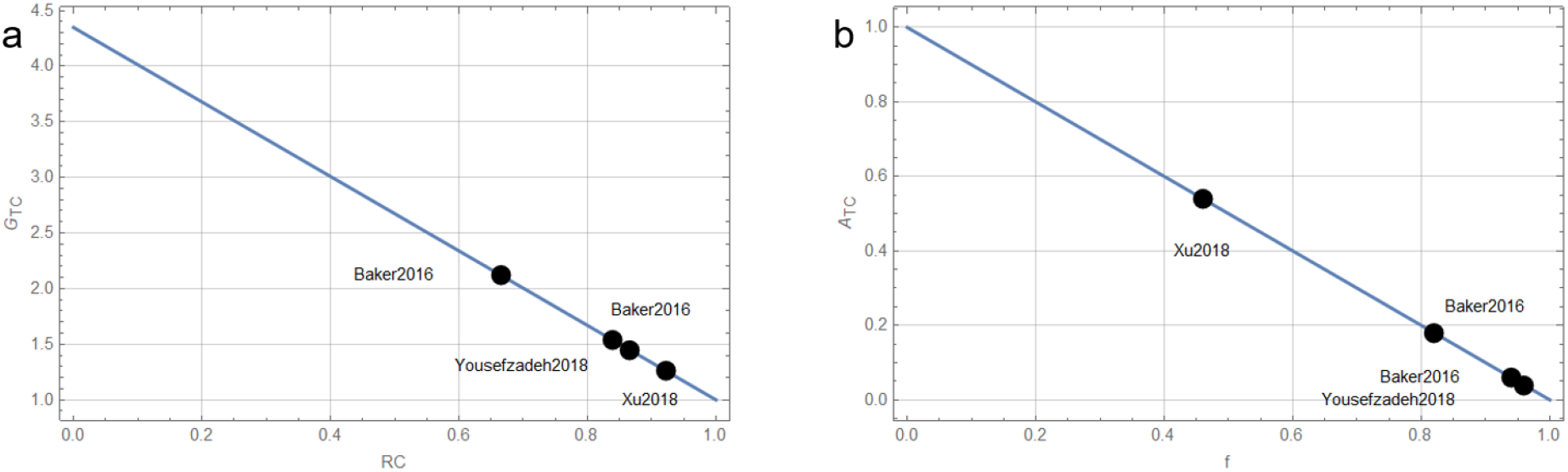
(a) Analysis of scenario C showing by how much the repair capacity has to be affected by the presence of senolytic drugs to result in the *G*_*TC*_ values observed experimentally and (b) which fraction of all age-related damage types has to be sensitive to senolytic treatment to give the *A*_*TC*_ values observed experimentally. Surprisingly, for some studies this value has to be around 95%.

Parameter *A* is equivalent to the mortality immediately after the start of treatment. In our simulations mortality is directly proportional to the combined damage of class 1 (*D*_1_) and class 2 (*D*_2_), which, until the first senolytic treatment, are given by

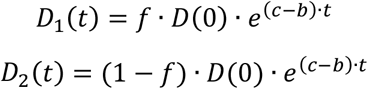

where *f* is the fraction of the total age-related damage that belongs to D_1_. Since repeated senolytic treatments effectively remove all class 1 damage it follows that:

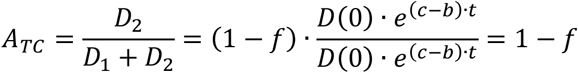

Fig. 10 presents these results graphically by showing how much *RC* has to be reduced so that Scenario C yields *G*_TC_ values observed in the experimental studies (a) and what fraction of all damage types have to respond to senolytic treatment (class 1) to obtain the *A*_TC_ values observed experimentally (b). *RC* has to be reduced between 8% and 34% (0.08 < *hRC* < 0.34) to be compatible with the studies, but the specific numerical values depend on the fraction of new age-related damage that is normally removed by the repair system (*b*/*c*), here 0.154/0.2=0.77. The larger this fraction, the smaller is the impairment that is required to be consistent with experimental observation. Unfortunately, this fraction is currently unknown and therefore represents a free parameter of Scenario C. The situation is somewhat different for *f*, the fraction of all age-related damage that responds to senolytic treatment. To be consistent with experiments *f* has to be between 46% and 96%, which is remarkably large and which does not rely on other unknown model parameters. But this finding is in line with recent experimental findings that senescent cells directly or via the SASP cause or promote a bewildering array of age-related damage and diseases ranging from osteoarthritis (Xu *et al.*, 2017), hepatic steatosis (Ogrodnik *et al.*, 2017) and senile lentigo (Yoon *et al.*, 2018) to type 1 diabetes (Thompson *et al.*, 2019), cognitive impairment (Ogrodnik *et al.*, 2021) and different types of cancer (Wang *et al.*, 2017; Yanai & Fraifeld, 2018).

If we consider the strengths and limitations of this study, the immediate strengths are the detailed examination of the post-senolytic impacts on compression of mortality (in particular, revealing the increase in actuarial ageing rate *G*) and the preliminary exploration via modelling of how these impacts might be explained mechanistically. There are of course limitations in the models. Firstly, in the absence of experimental evidence, we have had to make quite simple assumptions in building Scenarios A, B and C. Nevertheless, we believe the scenarios to capture the most important possibilities that need to be explored, providing a basis for further development. Secondly, we have needed to set parameter values rather arbitrarily. We have not undertaken exhaustive exploration of the parameter combinations that could affect the outcomes, but we believe the reported results to be valuable, if only as proof-of-principle. Thirdly, we have confined the stochastic elements of the model to the random selection, according to mortality risk, of individual deaths. In theory, the deterministic equations governing damage accumulation in the three scenarios could also be made stochastic. However, there is no basis in data to introduce the extra mathematical complexity and increased number of parameters that would be involved, and it seems unlikely that the exploratory potential of the models, for the purposes we have pursued, would be thereby enhanced.

If our main assumption of Scenario C, that repeated treatments with senolytics impair the organism’s repair capacity, is correct, then a single senolytic treatment, as shown in Fig. 9, might represent an optimized treatment schedule. Since the repair system can recover from the single treatment, it would not lead to an accelerated accumulation of senolytics resistant damage and consequently also maximum lifespan should be increased. A survival curve of such a population should also not exhibit an elevated parameter *G* as seen in the publications that we investigated here.

In conclusion, the results from senolytic experiments present intriguing challenges in terms both of understanding the effects of senescent cells on the ageing process and of potentially delivering interventions to improve healthy ageing in humans. The post-senolytic compression of mortality in mice might be ultimately beneficial but as we have revealed comes at the cost of an increased actuarial ageing rate. Since this suggests the presence of side effects associated with the senolytic treatment, it is important to consider what these may be. In line with our recent examination of the evolutionary basis of cellular senescence (Kowald *et al.*, 2020), we suggest that senolytic treatments, especially when repeated, might impair the repair capacity delivered for example by the surveillance of the immune system. If this is correct, it suggests ways to optimise the administration of senolytic therapies

## CRediT authorship contribution statement

Conceptualization: A.K., T.K.

Writing review and editing: A.K., T.K.

Visualization: A.K.

Coding: A.K.

## Acknowledgements

AK acknowledges financial support by the Federal Ministry of Education and Research (BMBF) of Germany for the SASKit study (FKZ 01ZX103A). TK acknowledges support from the NORDEA Foundation (02-2013-0220), Denmark, and Novo Nordisk Foundation (NNF17OC00278), Denmark.

